# Impacts of *perR* on oxygen sensitivity, gene expression, and murine infection in *Clostridioides difficile* 630Δ*erm*

**DOI:** 10.1101/2024.10.30.621113

**Authors:** Anna L. Gregory, Hailey E. Bussan, Madeline A. Topf, Andrew J. Hryckowian

## Abstract

*Clostridioides difficile* infection (CDI), characterized by colitis and diarrhea, afflicts approximately half a million people in the United States every year, burdening both individuals and the healthcare system. *C. difficile* 630Δ*erm* is an erythromycin-sensitive variant of the clinical isolate *C. difficile* 630 and is commonly used in the *C. difficile* research community due to its genetic tractability. 630Δ*erm* possesses a point mutation in *perR*, an autoregulated transcriptional repressor that regulates oxidative stress resistance genes. This point mutation results in a constitutively de-repressed PerR operon in 630Δ*erm*. To address the impacts of *perR* on phenotypes relevant for oxygen tolerance and relevant to a murine model of CDI, we corrected the point mutant to restore PerR function in 630Δ*erm* (herein, 630Δ*erm perR*^WT^). We demonstrate that there is no difference in growth between 630Δ*erm* and a 630Δ*erm perR*^WT^ under anaerobic conditions or when exposed to concentrations of O_2_ that mimic those found near the surface of the colonic epithelium. However, 630Δ*erm perR*^WT^ is more sensitive to ambient oxygen than 630Δ*erm*, which coincides with alterations in expression of a variety of *perR*-dependent and *perR*-independent genes. Finally, we show that 630Δ*erm* and 630Δ*erm perR*^WT^ do not differ in their ability to infect and cause disease in a well-established murine model of CDI. Together, these data support the hypothesis that the *perR* mutation in 630Δ*erm* arose as a result of exposure to ambient oxygen and that the *perR* mutation in 630Δ*erm* is unlikely to impact CDI-relevant phenotypes in laboratory studies.

**Importance:** *Clostridioides difficile* is a diarrheal pathogen and a major public health concern. To improve humans’ understanding of *C. difficile*, a variety of *C. difficile* isolates are used in research, including *C. difficile* 630Δ*erm*. 630Δ*erm* is a derivative of the clinical isolate 630 and is commonly studied because it is genetically manipulable. Previous work showed that a mutation in *perR* in 630Δ*erm* results in a dysregulated oxidative stress response, but no work has been done to characterize *perR*-dependent effects on the transcriptome or to determine impacts of *perR* during infection. Here, we identify transcriptomic differences between 630Δ*erm and* 630Δ*erm perR*^WT^ exposed to ambient oxygen and demonstrate that there is no strain-based difference in burdens in murine *C. difficile* infection.

## Introduction

*Clostridioides difficile* is a leading cause of infectious diarrhea, resulting in an estimated 500,000 annual cases in the US alone (1). A healthy microbiota typically prevents symptomatic *C. difficile* infection (CDI) through colonization resistance. However, ecological disturbances, commonly broad-spectrum antibiotics, disrupt the gut microbiota and give *C. difficile* access to vacant niches, which facilitates CDI (2–4). During dysbiosis, *C. difficile* can access nutrients otherwise consumed by the gut microbiota, overcome the effects of inhibitory metabolites produced by the microbiota/host, and increase in abundance in the gastrointestinal tract. During infection, *C. difficile* produces toxins (TcdA and TcdB; and CDT in hypervirulent strains) which induce host inflammation and are responsible for the diarrhea characteristic of CDI. This diarrhea contributes to transmission of *C. difficile* spores and allows *C. difficile* to gain a metabolic advantage by suppressing the recovery of the microbiota (3, 5–7). However, by inducing inflammation, *C. difficile* also causes the elevation of oxygen (O_2_) and reactive oxygen species (ROS) in the gut lumen (8–12). As an obligate anaerobe, *C. difficile* has evolved a variety of strategies to resist oxidative stress including sporulation, a versatile metabolism, and oxygen detoxification enzymes such as flavodiirons, rubrerythrins, and desulfoferrodoxin (9, 13–15). Previous work described various aspects of the response of *C. difficile* to oxidative stress (8, 16–18) and characterized proteins that play roles in oxidative stress tolerance (8–10, 16–19). One important regulator of a subset of these oxygen detoxification enzymes is the peroxide repressor (PerR).

PerR is an autoregulated transcriptional repressor and is a member of the ferric uptake repressor (Fur) family of proteins (19, 20). PerR is involved in oxidative stress responses in multiple bacterial species including *Clostridium acetobutylcium, Bacillus subtilis, Streptococcus pyogenes, Streptococcus mutans, Staphylococcus aureus, Campylobacter jejuni,* and *Clostridioides difficile* (21–27). Under anaerobic conditions, PerR binds to its target promoters and represses their expression. Oxidative stress triggers a metal catalyzed histidine oxidation and PerR undergoes a conformational change, causing it to release from its target promoters, which induces de-repression of the PerR regulon (20, 27). In *S. pyogenes,* a *perR* mutant was hyper-resistant to peroxide. However, it was highly attenuated in a murine model, demonstrating the importance of an appropriately regulated PerR regulon for virulence *in vivo* (22). A *perR* mutation identified in a *S. mutans* Δ*spxA1* strain similarly rendered PerR inactive, priming the strain to tolerate oxidative stress. However, this work also showed that PerR had a limited impact on the transcriptional response of *S. mutans* to hydrogen peroxide (24). In *C. acetobutylcium*, an obligate anaerobe, a *perR* mutant was more aerotolerant than wildtype and it was determined that PerR regulates oxidative stress genes, including reverse rubrerythrins, flavordiiron proteins (FDPs), and superoxide-reducing desulfoferrodoxin (Dfx), as well as to two putative enzymes involved in central energy metabolism (28, 29).

In *C. difficile,* the operon containing *perR* consists of three genes: rubrerythrin (*rbr1*), *perR*, and a desulfoferrodoxin (*rbo*) (**Figure 1A**). Genes within this operon are upregulated upon exposure to 1.5% O_2_ *in vitro* (8). In ex-germfree mice mono-colonized with *C. difficile* and bi-colonized with *C. difficile* and *Bacteroides thetaiotaomicron,* the genes of the *perR* operon were among the 10% most abundant transcripts in *C. difficile* (2). Studies examining the activity of *rbo* demonstrated that when inactivated, *C. difficile* was more sensitive to oxygen exposure. Furthermore, when *C. difficile rbo* was expressed in *E. coli* it demonstrated superoxide scavenging activity (15). Taken together, these data indicate that *perR* expression is responsive to oxidative stress and that PerR and PerR-regulated genes are important for *C. difficile* to navigate oxidative stress both *in vitro* and perhaps *in vivo*.

**Figure 1:**
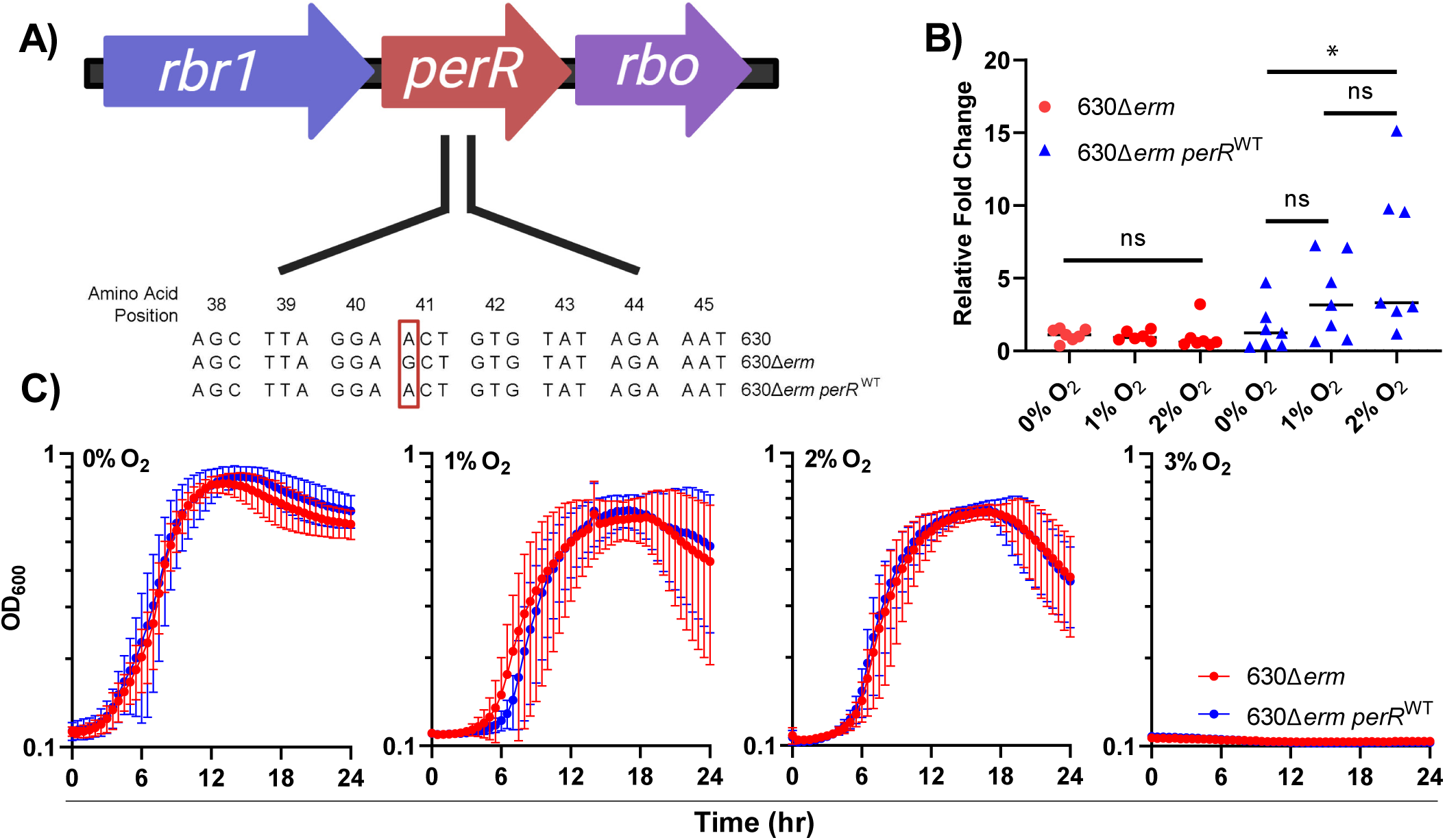
*In vitro* assays of 630Δ*erm* and 630Δ*erm perR*^WT^ exposed to physiologically-relevant levels of O_2_. (A) Genomic context of *perR* in *C. difficile* 630, 630Δ*erm* and 630Δ*erm perR*^WT^. The operon containing *perR* consists of three genes: a rubrerythrin (*rbr1*), the transcriptional repressor PerR (*perR*), and a desulfoferrodoxin (*rbo*). 630Δ*erm* has a point mutation in *perR*, resulting in a T41A amino acid substitution. (B) RT-qPCR of *perR* exposed to 0, 1, or 2% O_2_. Data points represent the relative fold change compared within each strain grown under anaerobic conditions, with *rpoC* used as the housekeeping gene (*n* = 7). Statistical testing was determined by Mann-Whitney test; *, *P* < 0.05. (C) Growth curves of 630Δ*erm* (red), and 630Δ*erm perR*^WT^ (blue) in mRCM grown anaerobically or hypoxically in the presence of 1, 2, and 3% O_2_. Data points represent the mean OD_600_ (*n* = 6-8) and error bars represent the standard deviation. Panel A was created with Biorender.com under agreement #Z41E368.

*C. difficile* 630Δ*erm* is an erythromycin-sensitive, lab-generated derivative of *C. difficile* 630 (herein, 630Δ*erm* and 630, respectively). 630Δ*erm* is amenable to allelic exchange procedures and is therefore commonly used in the field for generating knockout mutants. Seven spontaneous mutations were previously identified in 630Δ*erm* relative to 630. These mutations include a single nucleotide polymorphism (SNP) in *perR,* a SNP in *eutG,* a SNP in a transcriptional regulator of the GntR family (CD630_35630), and 3 SNPs in intergenic regions. Additionally, an 18 bp duplication is present in *spo0A* in 630Δ*erm,* which causes reduced sporulation efficiency compared to 630 (30). The *perR* point mutation in 630Δ*erm* results in an amino acid substitution at position 41 (T41A) (**Figure 1A**). This mutation affects the helix-turn-helix motif of the DNA binding domain of PerR and results in a constitutively expressed *perR* operon regardless of O_2_ or H_2_O_2_ exposure. A constitutively expressed *perR* operon provides 630Δ*erm* with a higher tolerance to O_2_ and H_2_O_2_ than parental strain 630 (27).

Despite these previous findings on the role of *perR* in *C. difficile* and other organisms, there are several remaining questions relating to the direct effects of the *perR* point mutation in 630Δ*erm* on oxidative stress resistance, gene expression, and the resulting impacts on infection. These are important gaps in knowledge, since this point mutation may impact interpretation of previous and future data since 630Δ*erm* is such a widely used strain in the *C. difficile* field. To address these gaps, we corrected the *perR* point mutation in 630Δ*erm* to create 630Δ*erm perR*^WT^. We demonstrate that this strain has a repressible *perR* operon and that there is no growth difference between 630Δ*erm* and 630Δ*erm perR*^WT^ in the presence of 0-3% O_2_. However, we show that 630Δ*erm* is fitter than 630Δ*erm perR*^WT^ when exposed to ambient air (21% O_2_). We also characterize 630Δ*erm* and 630Δ*erm perR*^WT^ transcriptomes exposed to ambient O_2_. Finally, using 630Δ*erm* and 630Δ*erm perR*^WT^, we demonstrate that functional PerR does not impact *C. difficile* burdens or diarrhea in a murine model of CDI.

## Results

### Targeted restoration of the mutant *perR* allele in 630Δ*erm* and impacts on growth and survival in the presence of oxygen

To determine the impact of a constitutively expressed PerR on *in vitro* phenotypes, we corrected the point mutation in 630Δ*erm*, generating 630Δ*erm perR*^WT^ (**Figure 1A**) (27). Correction of the point mutation was confirmed by whole-genome sequencing and alignment, comparing the parental 630Δ*erm* to 630Δ*erm perR*^WT^. A single SNP at position 1,006,274 (G → A) was identified, indicating the only genetic difference between the two strains was the restoration of a wild type *perR*.

To confirm wild type PerR function in 630Δ*erm perR*^WT^, *perR*-specific RT-qPCR was done on RNA extracted from 630Δ*erm* and 630Δ*erm perR*^WT^ grown in the presence of 0, 1, and 2% O_2_ (**Figure 1B**). These O_2_ concentrations were selected because they reflect those present in the colon (31, 32). This analysis revealed that *perR* transcription was non-responsive to oxygen in 630Δ*erm,* due to constitutive de-repression of its operon. However, *perR* transcripts in 630Δ*erm perR*^WT^ were elevated as a function of increasing oxygen exposure, which demonstrates the *perR* operon is de-repressed upon exposure to 2% O_2_. These data confirm that correcting the *perR* point mutation restored a wildtype, oxygen-responsive, phenotype and are supported by previously-published RNA-seq data that showed that *perR* is up-regulated in 630 at 1.5% O_2_ (8).

To evaluate the impacts of oxygen-responsive *perR* expression on *C. difficile* growth, 630Δ*erm* and 630Δ*erm perR*^WT^ were grown in a complex, rich medium (modified Reinforced Clostridial Medium [mRCM]) in the presence of 0, 1, 2, and 3% O_2_ (**Figure 1C & S1**) (3, 33). We did not observe differences in growth kinetics between the two strains at these O_2_ concentrations. However, growth of both strains was negatively impacted by increasing O_2_ concentration and no growth was observed in 3% O_2_. These data partially replicate results of a recent study on O_2_ reductases in *C. difficile*. There, targeted restoration of PerR function in 630Δ*erm* did not impact *C. difficile* growth at 1% O_2_ (34).

Despite differences in PerR activity between 630Δ*erm* and 630Δ*erm perR*^WT^, a constitutively derepressed PerR regulon offers no apparent fitness advantage at physiologically relevant O_2_ concentrations (**Figure 1**). Previous work showed that 630Δ*erm* has increased survival relative to 630 upon exposure to ambient O_2_ (27). To determine if 630Δ*erm perR*^WT^ has restored sensitivity an ambient air (∼21% O_2_), an ambient air exposure assay was performed. Cell viability of 630, 630Δ*erm,* and 630Δ*erm perR*^WT^ was quantified after exposure to ambient air for 0 to 90 minutes. Viability of 630 and 630Δ*erm perR*^WT^ were decreased upon exposure to ambient air. However, this exposure did not have an impact on 630Δ*erm,* demonstrating that targeted restoration of *perR* restores wild type levels of sensitivity to ambient air in *C. difficile* (**Figure 2**).

**Figure 2:**
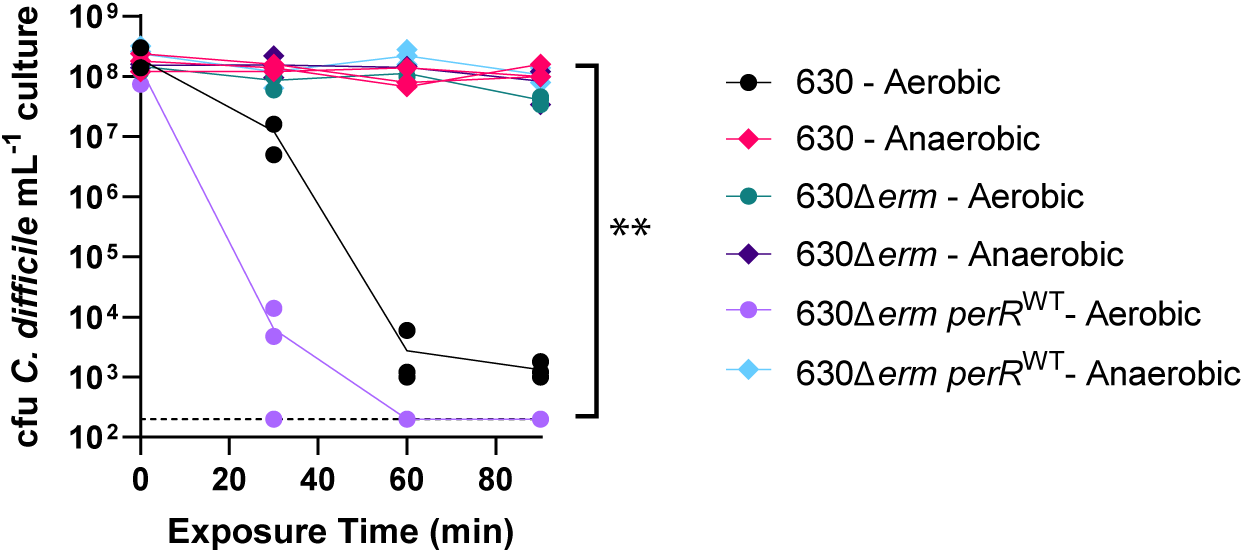
Aerotolerance assay of 630, 630Δ*erm,* and 630Δ*erm perR*^WT^ exposed to ambient air. Aliquots of stationary phase cultures of each strain were exposed to ambient air for 0, 30, 60, and 90 minutes or maintained under anaerobic conditions and plated on pre-reduced CDMN agar at the indicated time points. Colonies were quantified after overnight growth in an anaerobic chamber (*n* = 3 cultures per strain per condition). Statistical testing was determined by two-way ANOVA; **, *P* < 0.01.

### Differences in 630Δ*erm* and 630Δ*erm perR*^WT^ transcriptomes as a function of ambient air exposure

To better understand genes involved in 630Δ*erm* resistance to ambient air (**Figure 2**), we performed RNA-seq on 630Δ*erm* and 630Δ*erm perR*^WT^ at 0- and 60-minutes post ambient air exposure and compared transcriptomes between the two strains and two time points (**Tables S2-S5**). The 60-minute time point was chosen based on RT-qPCR of *perR* transcripts from total RNA extracted from 630 at 0-, 15-, 30-, and 60-minutes post air exposure (**Figure S2**), which showed elevated *perR* transcripts at both 30 and 60 minutes of aerobic exposure compared to anaerobic control. RNA extracted at 60 minutes was high-quality via Bioanalyzer (data not shown) and therefore selected as the timepoint. RNA-seq showed that the operon containing *perR* was responsive to aerobic exposure in 630Δ*erm perR*^WT^ but was expressed at high levels regardless of aerobic exposure in 630Δ*erm* (**Figure 3A**).

**Figure 3:**
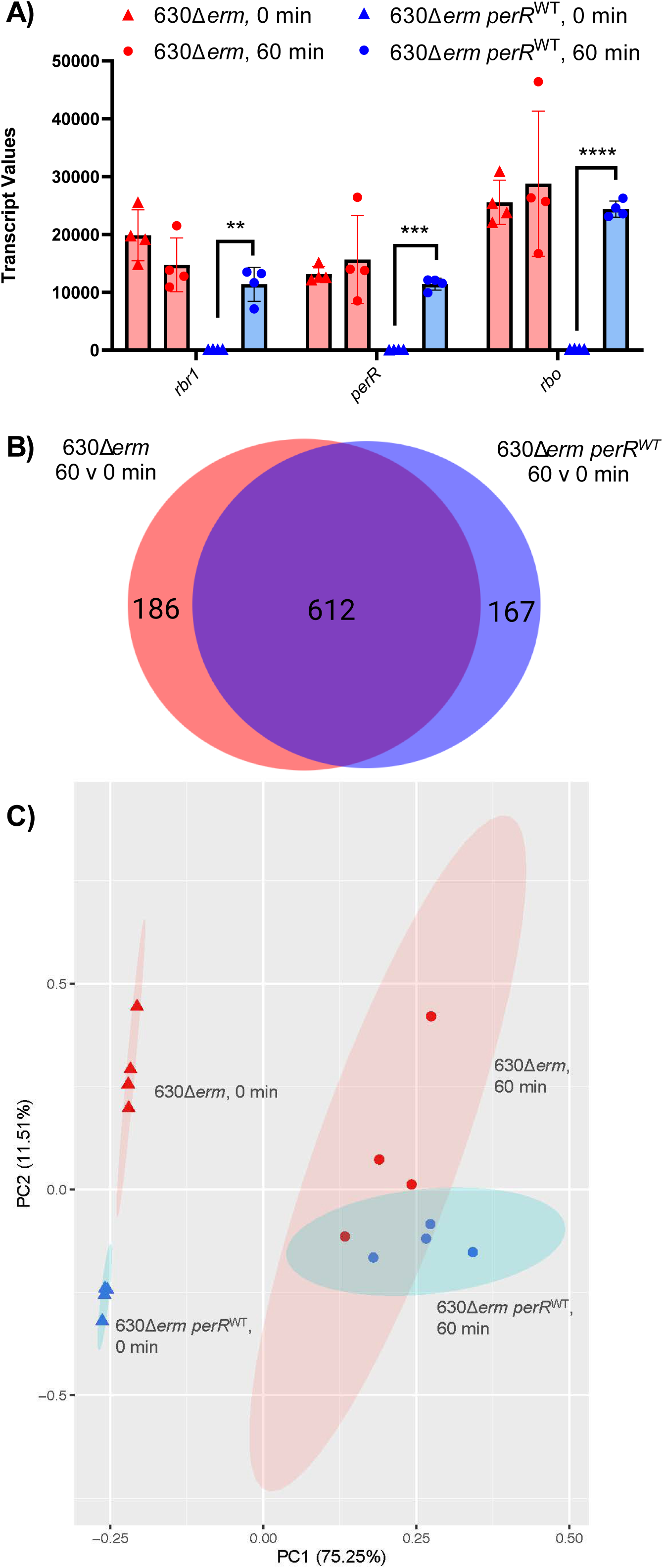
Transcriptional profiling of 630Δ*erm* and 630Δ*erm perR*^WT^ exposed to ambient air. 630Δ*erm* and 630Δ*erm perR*^WT^ were grown to mid-log phase in mRCM and then exposed to ambient air for 0 and 60 minutes. RNA-seq was performed on *n* = 4 independent cultures, for each strain at each timepoint. (A) Transcript values of the genes in the *perR* operon (*n* = 4). Statistical testing was determined by paired t-test; *, *P* < 0.05 **, *P* < 0.01, ***, *P* < 0.001, ****, *P* < 0.0001. (B) A Venn diagram illustrating that the majority of significantly differentially regulated genes (FC < |2|, *p* < 0.05) overlapped between the 630Δ*erm* and 630Δ*erm perR*^WT^ (C) Principal Component Analysis (PCA) plot of RNA-seq data from 630Δ*erm* (red) and 630Δ*erm perR*^WT^ (blue) exposed to ambient air for 0 (triangle) and 60 (circle) minutes. Ellipses represent 95% confidence intervals based on strain and time of ambient air exposure. Panel B was created with Biorender.com under agreement #N27K433.

Despite differences in expression of *rbr1, perR*, and *rbo* between the strains, there were largely overlapping patterns of gene expression for both strains between 0 and 60 minutes of ambient air exposure. Specifically, although there were differences in cell viability at the 60-minute time point (**Figure 2**), both strains shared 615 differentially regulated genes (**Figure 3B**), many of which were previously predicted/characterized to be involved in oxidative stress resistance (**Table S6**). This similarity in transcriptional response to ambient air is also evident from principal component analysis (**Figure 3C**), which shows that both strain and exposure to O_2_ had a significant impact on the transcriptome (the impact of time was more significant), but the interaction of the variables was not significant (PERMANOVA ∼ Time * Genotype, time p=0.001; strain p=0.006, time*strain, p = 0.085).

These data suggest that the increased viability of 630Δ*erm* relative to 630Δ*erm perR*^WT^ could be due to elevated levels of PerR-dependent gene products present in 630Δerm cells prior to oxygen exposure, which would prime the cells for oxidative stress. Beyond *rbr1, perR*, and *rbo,* 5 genes were up-regulated in 630Δ*erm* relative to 630Δ*erm perR*^WT^ at the t=0 timepoint. These genes include a putative oxidative stress glutamate dehydrogenase (CDIF630erm_00947), a putative metallo-beta-lactamase superfamily protein (CDIF630erm_00948), a putative conjugative transposon protein (CDIF630erm_01262), and two putative diguanylate kinase signaling proteins (CDIF630erm_00875 and CDIF630erm_01792). These data suggest that although PerR impacts *C. difficile* survival in ambient air, PerR does not exert major control, outside of the *perR* operon, over oxidative stress resistance genes in *C. difficile*, as previously observed in other microbes (21, 22, 24, 25, 28, 29).

### 630Δ*erm* and 630Δ*erm perR*^WT^ do not differ in their ability to infect or cause diarrhea in mice

During infection, *C. difficile* toxins induce host inflammation and elevate ROS during infection (35–37). Therefore, given that 630Δ*erm* and 630Δ*erm perR*^WT^ differ in their ability to survive oxidative stress, we sought to determine the impact of *perR* on strain fitness during infection. To examine this, we leveraged a well-established murine model of CDI (33, 38). Mice were placed on a fiber free diet and gavaged with clindamycin as in **Figure 4A** to reduce colonization resistance against *C. difficile*. Then, mice were gavaged with either 630Δ*erm* or 630Δ*erm perR*^WT^ to establish CDI. After 1 week of infection, the mice were switched to a high fiber diet to determine if fiber-dependent CDI clearance kinetics differ between the strains (38).

**Figure 4:**
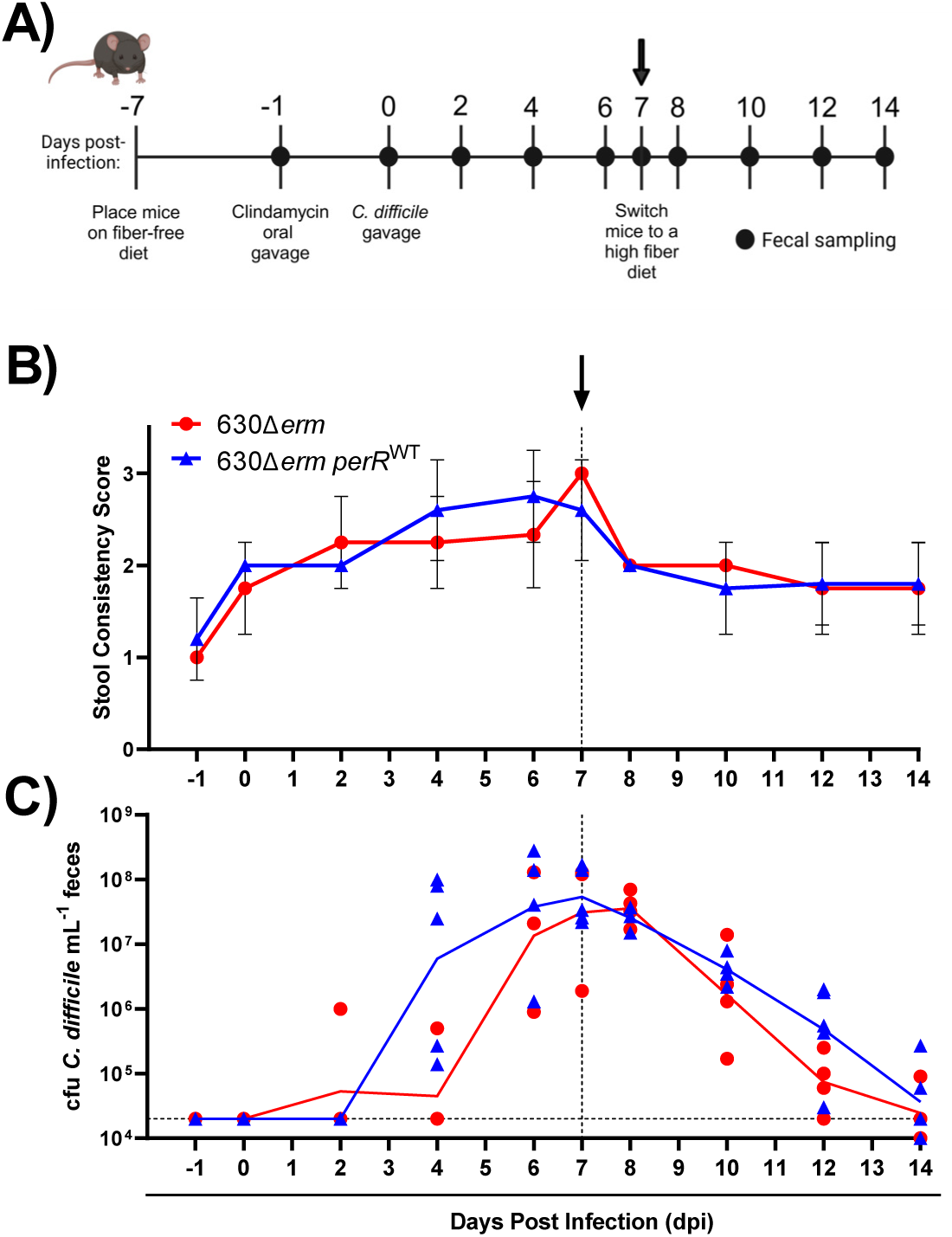
CDI in mice infected with 630Δ*erm* and 630Δ*erm perR*^WT^. (A) Conventional, age-matched male C57BL/6 mice were placed on a fiber-free diet, gavaged with clindamycin, and infected with either 630Δ*erm* or 630*Δerm perR*^WT^ (*n* = 4-5). Mice were switched to a high fiber diet seven days after being infected with *C. difficile*. (B) Average stool consistency scores: 1= hard, dry pellet; 2 = soft but fully formed pellet; 3 = runny, poorly formed pellet. (C) The geometric means of *C. difficile* burdens for each strain over the experimental time course. The limit of detection for this assay is indicated in the horizontal dashed line at 20,000 CFU *C. difficile*/mL feces. Panel A was created with Biorender.com under agreement #Z41E368. The vertical dashed line at seven days post infection in panels B and C indicates when the mice were switched from a fiber free to a high fiber diet.

There was no difference between the two strains in stool consistency scores nor *C. difficile* burdens across the 14 days (**Figure 4BC**). All mice experienced an increase in stool softness on 0 dpi due to administration of clindamycin (**Figure 4B**). However, after infection with *C. difficile,* mice experienced an increase in diarrhea, regardless of the strain with which they were infected. For both strains, fecal burdens of *C. difficile* were similar when on the fiber free diet, and decreased once the mice were placed on the high fiber diet, as previously demonstrated (**Figure 4C**) (38). Interestingly, there was a slight delay in *C. difficile* burdens in both strains at early time points. Specifically, consistent burdens were not seen until 4 dpi, possibly due to sensitivity of the strains to clindamycin (**Figure S3**). Taken together, these data demonstrate that 630Δ*erm* and 630Δ*erm perR*^WT^ do not differ in their ability to infect or cause diarrhea in mice.

## Discussion

630Δ*erm* has been an important strain for elucidating the physiology and pathogenesis of *C. difficile* because of its extensive use to generate mutants via allelic exchange procedures. However, there are several mutations in 630Δ*erm* relative to its parent strain (a clinical isolate) that may impact generalizability of the findings made in 630Δ*erm* to other *C. difficile* strains. The mutations in 630Δ*erm* include a point mutation in *perR* that renders its PerR regulon constitutively de-repressed. Despite knowledge of this *perR* point mutation and its impacts on expression of genes within the *perR* operon, the effects of this mutation on *C. difficile* oxidative stress resistance, on the broader *C. difficile* transcriptome, and on CDI phenotypes remained poorly characterized.

Here, we restored a wild type copy of *perR* in 630Δ*erm* to create 630Δ*erm perR^WT^*. We determined that there is no difference in growth between 630Δ*erm* and *630*Δ*erm perR^WT^*at physiologically-relevant O_2_ levels (**Figure 1**). However, using these strains, we showed that a constitutively de-repressed PerR regulon allows *C. difficile* to tolerate ambient air exposure (**Figure 2**). Previous work that compared the survival of *C. difficile* exposed to ambient O_2_ compared 630 and 630Δ*erm* which have multiple genetic differences in addition to the *perR* point mutation (27). Our results support the hypothesis that the differences in oxidative stress resistance in these strains was due to *perR* and not these other mutations. In addition, given that the point mutation in *perR* is unique to 630Δ*erm* when compared to 11 diverse clinical isolates of *C. difficile* (27), it is reasonable to assume that this mutation was selected for due to oxygen exposure during laboratory passage.

To better understand the genes repressed by PerR (which likely contribute to increased O_2_ tolerance by 630Δ*erm*), we performed RNA-seq on 630Δ*erm* and 630Δ*erm perR*^WT^ exposed to ambient air (**Figure 3**). This analysis suggested that PerR represses a small fraction of genes in *C. difficile*. Under anaerobic conditions, 8 genes were up-regulated in 630Δ*erm* relative to 630Δ*erm perR*^WT^ (**Table S2**). This includes the genes present in the *perR* operon (*rbr1, perR,* and *rbo*), an oxidative stress glutamate dehydrogenase, a putative metallo-beta-lactamase, a putative conjugative transposon protein, and two putative diguanylate kinase signaling proteins. Our RNA-seq data also show that large-scale changes to the *C. difficile* transcriptome occur at 60 minutes post air exposure, regardless of whether the strain has functional PerR. Specifically, 615 shared genes are differentially regulated in both strains at 60 minutes post-exposure to ambient air relative to the pre-exposure time point. This include genes involved in oxidative stress resistance (**Table S6**), many of which are likely under the control of other oxidative stress regulators (e.g. σ^B^) 12–14, 34, 42). Despite similarities of the transcriptional responses of 630Δ*erm* and 630Δ*erm perR*^WT^ to ambient air, 630Δ*erm* and 630Δ*erm perR*^WT^ have some differences in their responses to this treatment (**Figure 3**). Deeper analysis of functional categories of these genes (**Tables S6-S8**) revealed oxygen-dependent up-regulation of genes involved in Stickland fermentation, ribosome synthesis, and the CRISPR system in 630Δ*erm* and oxygen-dependent down-regulation of peptidoglycan and teichoic acid metabolism in 630Δ*erm perR*^WT^. These changes in gene expression mirror the differential survival of these two strains in the presence of oxygen (39).

Our data also indicate that there is no difference between 630Δ*erm* and *630*Δ*erm perR^WT^* in *C. difficile* burdens nor severity of infection in a murine model of CDI (**Figure 4**). While our work was done in conventional mice, previous work showed that *rbo, perR*, and *rbr* were among the top 10% most highly expressed genes in gnotobiotic mice infected with 630 (2), suggesting that PerR-dependent gene expression is important during infection. Because longitudinal and radial oxygen gradients are present in the gastrointestinal tract and O_2_ levels can be elevated by antibiotic treatment, it is possible that *C. difficile* encounters enough O_2_ to de-repress PerR-dependent genes during the onset, establishment, or maintenance of murine infection (31, 32, 40, 41). Therefore, the mouse experiments performed may disguise possible differences in fitness (positive or negative) due to a constitutively de-repressed PerR operon.

In summary, our work establishes that a constitutively de-repressed PerR regulon offers no fitness advantage at O_2_ levels encountered in the distal GI tract (**Figure 1**) nor in a mouse model of CDI (**Figure 4**). However, a constitutively de-repressed PerR regulon provides *C. difficile* with tolerance to ambient air (**Figure 2**) and impacts gene expression in *C. difficile* in the presence and absence of oxygen (**Figure 3**). Unique to 630Δ*erm*, the *perR* mutation invokes consideration of selective pressures that this strain may have encountered during exposure to ambient air in laboratory settings (27). This study adds to a growing body of literature on the ways in which obligate anaerobes resist oxidative stress and will contribute to future work in understanding these responses in *C. difficile*.

## Materials and Methods

### Bacterial strains and culture conditions

*C. difficile* strains 630, 630Δ*erm* and 630Δ*erm perR*^WT^ (42, 43) were maintained as ࢤ80°C stocks in 25% glycerol under anaerobic conditions in septum-topped vials. *C. difficile* strains were struck out on CDMN agar, or *C. difficile* agar base (Oxoid) supplemented with 32 mg/L moxalactam (Santa Cruz), 12 mg/L norfloxacin (Sigma Aldrich), and 7% defibrinated horse blood (HemoStat Laboratories), and cultured anaerobically for 24 hours. A single colony was picked into 5 mL of pre-reduced BD Difco^TM^ reinforced clostridial medium (RCM) or a modified RCM (mRCM; RCM without soluble starch and agar)(33). Liquid cultures were grown at 37°C, anaerobically for 16-24 hours and used as inocula for growth curves, aerotolerance assays, RNA-seq, RT-qPCR, and murine experiments. All bacterial growth media were pre-reduced for at least 24 hours in an anaerobic chamber (Coy) prior to use in experiments.

For *in vitro* growth curves, subcultures were prepared at 1:100 dilution in mRCM (3). Growth curves were performed anaerobically or in a hypoxic chamber at 1%, 2%, and 3% O_2_ (Coy). Clindamycin sensitivity growth curves were performed anaerobically in mRCM with clindamycin concentrations spiked into each well. All growth curves were performed in sterile polystyrene 96- well tissue culture plates (Falcon) with low evaporation lids using a BioTek Epoch2 plate reader at 30-minute intervals. Plates were shaken on the orbital setting for 10 seconds before each read. The OD_600_ of the cultures was recorded using Gen5 software (version 3.10.06).

### Generation of C. difficile 630Δerm perR^WT^

The *perR* point mutation was corrected using the PyrE allelic exchange system (44). Primers indicated in **Table S1** were used to amplify *perR* from 630 genomic DNA. The amplicon containing *perR* from 630 was ligated into pMTL-YN3 after AscI and SbfI digestion of the vector and inserted using New England Biolabs Quick Ligation Kit (M2200S). The plasmid construct was transformed and propagated into One Shot Top10 *E. coli* (Invitrogen) before transformation into conjugation proficient *E. coli* HB101/pRK24 cells. The pMTL-YN3 with 630 *perR* was conjugated into 630Δ*erm*Δ*pyrE*. Plasmid integrants were selected for using BHIS supplemented with thiamphenicol (10 or 15 μg/mL), cefoxitin (8 μg/mL), kanamycin (30 or 50 μg/mL), and uracil (5 μg/mL). Double crossover events were selected for using a defined minimal medium for *C. difficile* (CDDM) supplemented with uracil (5 μg/mL) and 5-fluoroorotic acid (2 mg/mL). Whole colony PCR using GoTaq Green Master Mix (Promega) amplified *perR* from potential clones. HypC4III digestion of PCR purified *perR* (Zymo DNA Clean and Concentrator-5) confirmed 630 *perR* integration since this restriction enzyme digests 630 *perR* but is unable to recognize that cut site in 630Δ*erm* due to the point mutation (**Figure S4**). After confirmation of 630 *perR* integration, the *pyrE* locus was restored using pMTL-YN1C to generate 630Δ*erm perR*^WT^.

Illumina whole genome sequencing on DNA extracted (45) from 630Δ*erm* and 630Δ*erm perR*^WT^ was performed by Microbial Genome Sequencing Center (MiGS) with a minimum read count of 1.33 million reads (200 Mbp) per sample. Raw sequencing data were assembled using a reference-guided assembly pipeline (https://github.com/pepperell-lab/RGAPepPipe_CHTC) to *C. difficile* strain 630Δ*erm* (GenBank: LN614756.1) as previously described (46). Briefly, Fastqc v0.12.1 (47) assessed the quality of the sequences, which were then trimmed using Trimmomatic v0.39 (48). Sequences were aligned to the reference using BWA Mem v0.7.18 (49), and alignments were processed using SAMtools v1.21 (50). Picard v 2.18.25 (https://github.com/broadinstitute/picard) was used to remove duplicates and add read groups. Pilon v 1.78 (51) identified variants. Assembly and alignment quality was determined using Qualimap BamQC v2.2.1 (52). The mean coverage was 160X for 630Δ*erm* and 50X for 630Δ*erm perR*^WT^. Both samples had >95% aligned reads to the reference. To identify the SNP difference between 630Δ*erm* and 630Δ*erm perR*^WT^, a VCF was created using SnpSites v2.5.1 (53).

### Aerotolerance assays

Liquid cultures of 630, 630Δ*erm* and 630Δ*erm perR*^WT^ were grown anaerobically in RCM overnight. 200 μL of each culture was aliquoted into sterile polystyrene 96-well tissue culture plates, with four replicate plates set up, and exposed to ambient air for 0, 30, 60 and 90 minutes at room temperature. Aeration of cultures using a multichannel pipette was performed immediately after removing from the chamber and approximately every 10 minutes throughout the assay. At each timepoint, one of the 96-well plates was passaged back into the anaerobic chamber. Serial dilutions were performed using pre-reduced PBS and plated on pre-reduced CDMN. Plates were incubated anaerobically for 24 hours at 37°C, and colonies present on CDMN plates were quantified.

### Transcriptional profiling of *C. difficile* response to ambient air

*C. difficile* 630Δ*erm* and 630Δerm *perR*^WT^ overnight cultures were back-diluted 1:100 into 35 mL of pre-reduced mRCM in Erlenmeyer flasks and incubated anaerobically at 37°C until cultures reached mid-log phase (OD_600_=0.3-0.4). At mid-log phase, 5 mL aliquots of the cultures were diluted 1:1 in chilled 1:1 ethanol:acetone and stored at ࢤ20° to preserve RNA. The remaining cultures were aerobically shaken (220 rpms) at 37°C for 60 minutes. After 60 minutes, 5 mL aliquots were diluted 1:1 in chilled 1:1 ethanol:acetone and stored at ࢤ20° to preserve RNA (54).

RNA was extracted by centrifuging samples at 3,000 xg for 5 minutes at 4°C. Pellets were washed with 5 mL cold, nuclease free PBS, and centrifuged at 3,000 xg for 5 minutes at 4°C. The supernatant was removed, and remaining pellets were resuspended in 1 mL TRIzol and processed using a TRIzol Plus RNA Purification Kit (Thermo) with on-column DNase treatment. Purified RNA integrity was confirmed via 2100 Agilent BioAnalyzer and frozen at ࢤ80°C.

RNA-seq was performed by Microbial Genome Sequencing Center (MiGS) on high-quality rRNA-depleted RNA extracts (12 million paired end reads per sample). Quality control and adapter trimming was performed with bcl2fastq (version 2.20.0.445)(55). Read mapping was performed with HISAT2 (version 2.2.0)(56). Read quantification was performed using Subread’s featureCounts (version 2.0.1)(57) functionality. Read counts loaded into R (version 4.0.2)(58) and were normalized using edgeR’s (59). Trimmed Mean of M values (TMM) algorithm (version 1.14.5). Subsequent values were then converted to counts per million (cpm). Differential expression analysis was performed using edgeR’s exact test for differences between two groups of negative-binomial counts with an estimated dispersion value of .1. Transcript level quantification, count normalization, and differential expression analysis were provided using *C. difficile* 630Δ*erm* (GCA_002080065.1_ASM208006v1) as the reference genome.

The Principal Coordinate Analysis (PCA) plot was generated from the RNA-seq data in R using version 4.4.1. PCA was performed using *prcomp* from *stats* package (version 4.4.1), and visualized using *ggplot2* (version 3.5.1). Confidence ellipses were generated using *stat_ellipse* as implemented in *ggplot2*. Permutation multivariate analysis of variance (PERMANOVA) was assessed by *vegan::adonis2* (version 2.6-6.1).

### RT-qPCR of *perR* in *C. difficile* exposed to varying levels of O_2_

To identify a timepoint for RNA-seq at which *perR* was de-repressed, *C. difficile* 630 overnight cultures were back-diluted 1:100 into 35 mL of pre-reduced RCM and incubated at 37°C until it reached mid-log phase (OD_600_=0.3-0.4). At mid-log phase, one set of cultures was incubated aerobically 37°C, shaking at 220 rpms, while the other culture was incubated at 37°C anaerobically. After 0, 15, 30, and 60 minutes, 5 mL aliquots of the cultures were diluted 1:1 in chilled 1:1 ethanol:acetone and stored at ࢤ20° to preserve RNA. RNA was extracted as described above.

RT-qPCR was performed using GoTaq 1-Step RT-qPCR Master Mix (Promega) according to the manufacturer’s instructions with a concentration of 1 ng/µL RNA in a final volume of 10 µL. Each reaction was run with three technical replicates. The RT-qPCRs were performed on a QuantStudio 7 Flex (Applied Biosystems) and threshold cycle (Ct) was determined using QuantStudio Real-Time PCR Software v1.7.2. The cycle run involved a reverse transcriptase activation and inactivation step of 40°C for 15 minutes and 95°C for 10 minutes. The PCR cycle was 95°C for 10 sec, 60°C for 30 sec and 72°C for 30 sec for 40 cycles, with a melt curve performed afterwards. Relative fold change was determined by comparing the Ct values against the average Ct value at 0 minutes for each condition.

To quantify *perR* transcript levels of *C. difficile* 630Δ*erm* and *C. difficile* 630Δ*erm perR*^WT^ exposed to low levels of O_2_, overnight cultures of each strain were back diluted 1:50 into 14 mL of pre-reduced mRCM in 100x15mm circular petri dishes (VWR) to optimize exposure to oxygen. Cultures were incubated at 37°C in 0, 1, and 2% O_2_ until they reached mid-log phase (OD_600_=0.3-0.4). At mid-log phase, 5 mL aliquots of the cultures were diluted 1:1 in chilled 1:1 ethanol:acetone and stored at ࢤ20° to preserve RNA and RNA was extracted as described above. RT-qPCR was performed using GoTaq 1-Step RT-qPCR Master Mix (Promega), with a concentration of 2 ng/μL of RNA in a final volume of 20 μL. Each reaction was set up with primers amplifying *perR* or *rpoC* (housekeeping gene) (**Table S1**), with 2-3 technical replicates. The RT-qPCRs were performed on a QuantStudio 7 Flex (Applied Biosystems) as previously described. Relative fold change was determined using the 2^-ΔΔCT^ method by comparing the Ct values to the anaerobic Ct value within each strain (60).

### Murine model of *C. difficile* infection (CDI)

All animal studies were done in strict accordance with the University of Wisconsin-Madison Institutional Animal Care and Use Committee (IACUC) guidelines (Protocol #M006305). CDI murine model was performed on age- and sex-matched, conventionally reared C57BL/6 inhouse between 6 and 9 weeks of age. Mice were fed a fiber free (FF) diet (Inotiv TD.150689) one week before antibiotic exposure. Mice were given a single dose of clindamycin by oral gavage (1 mg/mouse; 200 μL of 5-mg/mL solution), and 24 hours later were given 200 μL 630Δ*erm* or 630Δ*erm perR*^WT^ overnight cultures grown in RCM via oral gavage (*n* = 4-5 mice per condition; average inoculum 8.6×10^7^ CFU/mL).

At 7 dpi, mice were switched from the FF diet to a fiber rich standard rodent chow (Inotiv Teklad™ 2916) to observe fiber-dependent *C. difficile* clearance kinetics (33, 38). Throughout the entire experiment, feces were collected daily from mice directly into a microcentrifuge tube and kept on ice. To quantify *C. difficile* burdens, 1 μL of each fecal sample was collected with a disposable inoculating loop and resuspended in 200 μL PBS. 10-fold serial dilutions of fecal suspension were prepared in sterile polystyrene 96-well tissue culture plates (Falcon). For each sample, 10 μL aliquots of each dilution, with two technical replicates, were spread on CDMN agar. CDMN plates were incubated anaerobically at 37°C for 16-24 hours. Colonies were quantified and technical replicates were averaged to determine *C. difficile* burdens (limit of detection = 2 x 10^4^ CFU/mL). Stool consistency scores were also noted while processing fecal pellets. Fecal pellets were assigned a score of 1 = hard, dry pellets, difficult to transect with a disposable plastic culture loop, 2 = soft, fully formed pellets, easy to transect with a culture loop. or 3 = runny, poorly formed pellets, no pressure required to transect with a culture loop (3).

### Data availability

Data on normalized transcript abundance and differential expression analysis are found in **Tables S2-S5**. Prior to publication of a peer-reviewed manuscript, the raw data from the RNA-seq experiments shown in **Figure 3** and **Tables S2-S5** will be available from the corresponding author upon request. Similarly, raw data from whole genome sequencing of 630Δ*erm* and 630Δ*erm perR*^WT^ will be available from the corresponding author upon request prior to publication of a peer reviewed manuscript. These raw data will be uploaded to NCBI (Gene Expression Omnibus and Sequence Read Archive, respectively) and made freely available upon acceptance of the peer-reviewed manuscript.

### Statistical Analysis

All statistical analyses, except for the PERMANOVA, were performed using GraphPad Prism 9.4.1. PERMANOVA was performed as described in “PCA Generation and RNA-seq Data Analysis” section. Details of specific statistical analyses indicated in figure legends. For all figures, \**P*< 0.05, \*\**P*<0.01, \*\*\**P*<0.001, \*\*\*\**P*<0.0001.

## Acknowledgements

We thank Daniel Pensinger for guidance and scientific insight throughout this project and for technical support with the mouse experiments. We thank Alex Dobrila for insights and suggestions on troubleshooting various aspects of this project. We thank Joie Ling for maintaining the mouse breeding colony from which this work benefited. We thank Elise Cowley for support and advice regarding RT-qPCR.

This material is based upon work supported by startup funding from the University of Wisconsin-Madison (A.J.H.), the NSF GRFP (DGE-2137424 to A.L.G.), the Computation and Informatics in Biology and Medicine Training Grant (5T15LM007359 to H.E.B), and R01AI139261 (M.A.T).

Support was also provided by the Graduate School and the Office of the Vice Chancellor for Research and Graduate Education at the University of Wisconsin-Madison with funding from the Wisconsin Alumni Research Foundation. Any opinions, findings, and conclusions or recommendations expressed in this material are those of the authors and do not necessarily reflect the views of the funders. A.J.H. is the Judy and Sal A. Troia Professor in Gastrointestinal Disease Research at the University of Wisconsin-Madison.

A.L.G. and A.J.H. conceptualized the project. A.L.G. performed all laboratory experiments. A.L.G., H.E.B., and M.A.T. performed bioinformatic and statistical analyses and created the display items. A.L.G. and A.J.H. wrote the manuscript. All authors edited and approved the manuscript prior to submission.

## Supplemental figure and table legends

**Figure S1: Growth curves of (A) 630Δ*erm* and (B) 630Δ*erm perR*^WT^ in grown in mRCM at 0, 1, 2, and 3% O_2_.** Data points represent the mean OD_600_ (*n* = 6-8 cultures per strain per condition). Error bars represent the standard deviation. Related to Figure 1.

**Figure S2: RT-qPCR of *perR* transcripts from 630.** 630 was maintained under anaerobic conditions or exposed to ambient air for 0, 15, 30, and 60 minutes (*n* = 6 per strain per condition per time point). The relative fold change was determined by comparing the CT value against the average CT at 0 minutes for each condition. Statistical significance was determined by paired t-test; *, *P* < 0.05. Related to Figure 3.

**Figure S3: Growth curves of 630 and 630Δ*erm* in the presence of clindamycin.** Strains were grown in mRCM containing 0, 2.0, and 62.5 µg/mL clindamycin. Data points represent the mean OD_600_ (*n* = 2 cultures per strain per condition) and error bars represent the standard deviation. Related to Figure 4.

**Figure S4: HypC4III digest of *perR*-containing PCR amplicons.** Amplicons were generated from 630 and 630Δ*erm*. Due to the point mutation in *perR* in 630Δ*erm*, HypC4III digests the amplicon from 630Δ*erm* differently than 630. This technique was implemented to screen clones to identify 630Δ*erm perR*^WT^ candidates. Related to Figure 1.

**Table S1: Strains, plasmids, and oligonucleotides used in this study.** Related to Figures 1, 2, 3, and 4.

**Table S2: RNA-seq pair-wise comparisons of *C. difficile* 630Δ*erm* and 630Δ*erm perR*^WT^ at 0 minutes in response to ambient air.** Related to Figure 3.

**Table S3: RNA-seq pair-wise comparisons of *C. difficile* 630Δ*erm* and 630Δ*erm perR*^WT^ at 60 minutes in response to ambient air.** Related to Figure 3.

**Table S4: RNA-seq pair-wise comparisons of *C. difficile* 630Δ*erm* at 60 and 0 minutes in response to ambient air.** Related to Figure 3.

**Table S5: RNA-seq pair-wise comparisons of *C. difficile* 630Δ*erm perR*^WT^ at 60 and 0 minutes in response to ambient air.** Related to Figure 3.

**Table S6: RNA-seq fold change data from a selection of genes associated with oxidative stress.** Related to Figure 3.

**Table S7: Gene modules enriched within the genes differentially expressed in 630Δ*erm* in response to ambient air exposure.** Related to Figure 3.

**Table S8: Gene modules enriched within the genes differentially expressed in and 630Δ*erm perR*^WT^ in response to ambient air exposure.** Related to Figure 3.

## References

1. Guh AY, Mu Y, Winston LG, Johnston H, Olson D, Farley MM, Wilson LE, Holzbauer SM, Phipps EC, Dumyati GK, Beldavs ZG, Kainer MA, Karlsson M, Gerding DN, McDonald LC. 2020. Trends in U.S. Burden of Clostridioides difficile Infection and Outcomes. New England Journal of Medicine 382:1320–1330.

2. Ferreyra JA, Wu KJ, Hryckowian AJ, Bouley DM, Weimer BC, Sonnenburg JL. 2014. Gut microbiota-produced succinate promotes C. difficile infection after antibiotic treatment or motility disturbance. Cell Host Microbe 16:770–777.

3. Battaglioli EJ, Hale VL, Chen J, Jeraldo P, Ruiz-Mojica C, Schmidt BA, Rekdal VM, Till LM, Huq L, Smits SA, Moor WJ, Jones-Hall Y, Smyrk T, Khanna S, Pardi DS, Grover M, Patel R, Chia N, Nelson H, Sonnenburg JL, Farrugia G, Kashyap PC. 2018. Clostridioides difficile uses amino acids associated with gut microbial dysbiosis in a subset of patients with diarrhea. Sci Transl Med 10:eaam7019.

4. Jenior ML, Leslie JL, Young VB, Schloss PD. 2017. Clostridium difficile Colonizes Alternative Nutrient Niches during Infection across Distinct Murine Gut Microbiomes 2:19.

5. Koenigsknecht MJ, Theriot CM, Bergin IL, Schumacher CA, Schloss PD, Young VB. 2015. Dynamics and Establishment of Clostridium difficile Infection in the Murine Gastrointestinal Tract. Infect Immun 83:934–941.

6. Jenior ML, Leslie JL, Young VB, Schloss PD. 2018. Clostridium difficile Alters the Structure and Metabolism of Distinct Cecal Microbiomes during Initial Infection To Promote Sustained Colonization. mSphere 3.

7. Litvak Y, Byndloss MX, Bäumler AJ. 2018. Colonocyte metabolism shapes the gut microbiota. Science 362:eaat9076.

8. Weiss A, Lopez CA, Beavers WN, Rodriguez J, Skaar EP. 2021. Clostridioides difficile strain-dependent and strain-independent adaptations to a microaerobic environment. Microb Genom 7.

9. Giordano N, Hastie JL, Smith AD, Foss ED, Gutierrez-Munoz DF, Carlson PE. 2018. Cysteine Desulfurase IscS2 Plays a Role in Oxygen Resistance in Clostridium difficile. Infection and Immunity 86:8.

10. Britton RA, Young VB. 2014. Role of the Intestinal Microbiota in Resistance to Colonization by *Clostridium difficile*. Gastroenterology 146:1547–1553.

11. Theriot CM, Young VB. 2015. Interactions Between the Gastrointestinal Microbiome and Clostridium difficile. Annu Rev Microbiol 69:445–461.

12. Kint N, Alves Feliciano C, Martins MC, Morvan C, Fernandes SF, Folgosa F, Dupuy B, Texeira M, Martin-Verstraete I. 2020. How the Anaerobic Enteropathogen Clostridioides difficile Tolerates Low O2 Tensions. mBio 11:e01559–20.

13. Kint N, Janoir C, Monot M, Hoys S, Soutourina O, Dupuy B, Martin-Verstraete I. 2017. The alternative sigma factor σB plays a crucial role in adaptive strategies of Clostridium difficile during gut infection. Environmental Microbiology 19:1933–1958.

14. Folgosa F, Martins MC, Teixeira M. 2018. The multidomain flavodiiron protein from Clostridium difficile 630 is an NADH:oxygen oxidoreductase. 1. Sci Rep 8:10164.

15. Kochanowsky R, Carothers K, Roxas BAP, Anwar F, Viswanathan VK, Vedantam G. 2024. Clostridioides difficile superoxide reductase mitigates oxygen sensitivity. Journal of Bacteriology 206:e00175–24.

16. Edwards AN, Karim ST, Pascual RA, Jowhar LM, Anderson SE, McBride SM. 2016. Chemical and Stress Resistances of Clostridium difficile Spores and Vegetative Cells. Front Microbiol 7.

17. Giordano N, Hastie JL, Carlson PE. 2018. Transcriptomic profiling of Clostridium difficile grown under microaerophillic conditions. Pathogens and Disease 76:fty010.

18. Neumann-Schaal M, Metzendorf NG, Troitzsch D, Nuss AM, Hofmann JD, Beckstette M, Dersch P, Otto A, Sievers S. 2018. Tracking gene expression and oxidative damage of O2-stressed Clostridioides difficile by a multi-omics approach. Anaerobe 53:94–107.

19. Duarte V, Latour J-M. 2013. PerR: a bacterial resistance regulator and can we target it? Future Med Chem 5:1177–1179.

20. Lee J-W, Helmann JD. 2006. The PerR transcription factor senses H2O2 by metal-catalysed histidine oxidation. Nature 440:363–367.

21. Bsat N, Herbig A, Casillas-Martinez L, Setlow P, Helmann JD. 1998. Bacillus subtilis contains multiple Fur homologues: identification of the iron uptake (Fur) and peroxide regulon (PerR) repressors. Mol Microbiol 29:189–198.

22. Brenot A, King KY, Caparon MG. 2005. The PerR regulon in peroxide resistance and virulence of Streptococcus pyogenes. Molecular Microbiology 55:221–234.

23. King KY, Horenstein JA, Caparon MG. 2000. Aerotolerance and peroxide resistance in peroxidase and PerR mutants of Streptococcus pyogenes. J Bacteriol 182:5290–5299.

24. Kajfasz JK, Zuber P, Ganguly T, Abranches J, Lemos JA. 2021. Increased Oxidative Stress Tolerance of a Spontaneously Occurring perR Gene Mutation in Streptococcus mutans UA159. Journal of Bacteriology 203:10.1128/jb.00535-20.

25. Horsburgh MJ, Clements MO, Crossley H, Ingham E, Foster SJ. 2001. PerR Controls Oxidative Stress Resistance and Iron Storage Proteins and Is Required for Virulence in Staphylococcus aureus. Infect Immun 69:3744–3754.

26. Kim M, Hwang S, Ryu S, Jeon B. 2011. Regulation of perR Expression by Iron and PerR in Campylobacter jejuni ▿. J Bacteriol 193:6171–6178.

27. Troitzsch D, Zhang H, Dittmann S, Düsterhöft D, Möller TA, Michel A-M, Jänsch L, Riedel K, Borrero-de Acuña JM, Jahn D, Sievers S. 2021. A Point Mutation in the Transcriptional Repressor PerR Results in a Constitutive Oxidative Stress Response in Clostridioides difficile 630Δerm. mSphere 6:e00091–21.

28. Hillmann F, Fischer R-J, Saint-Prix F, Girbal L, Bahl H. 2008. PerR acts as a switch for oxygen tolerance in the strict anaerobe Clostridium acetobutylicum. Mol Microbiol 68:848–860.

29. Hillmann F, Döring C, Riebe O, Ehrenreich A, Fischer R-J, Bahl H. 2009. The Role of PerR in O2-Affected Gene Expression of Clostridium acetobutylicum. J Bacteriol 191:6082– 6093.

30. Collery MM, Kuehne SA, McBride SM, Kelly ML, Monot M, Cockayne A, Dupuy B, Minton NP. 2017. What’s a SNP between friends: The influence of single nucleotide polymorphisms on virulence and phenotypes of Clostridium difficile strain 630 and derivatives. Virulence 8:767–781.

31. Singhal R, Shah YM. 2020. Oxygen battle in the gut: Hypoxia and hypoxia-inducible factors in metabolic and inflammatory responses in the intestine. J Biol Chem 295:10493– 10505.

32. He G, Shankar RA, Chzhan M, Samouilov A, Kuppusamy P, Zweier JL. 1999. Noninvasive measurement of anatomic structure and intraluminal oxygenation in the gastrointestinal tract of living mice with spatial and spectral EPR imaging. Proc Natl Acad Sci U S A 96:4586–4591.

33. Pensinger DA, Fisher AT, Dobrila HA, Van Treuren W, Gardner JO, Higginbottom SK, Carter MM, Schumann B, Bertozzi CR, Anikst V, Martin C, Robilotti EV, Chow JM, Buck RH, Tompkins LS, Sonnenburg JL, Hryckowian AJ. 2023. Butyrate Differentiates Permissiveness to Clostridioides difficile Infection and Influences Growth of Diverse C. difficile Isolates. Infection and Immunity 91:e00570–22.

34. Caulat LC, Lotoux A, Martins MC, Kint N, Anjou C, Teixeira M, Folgosa F, Morvan C, Martin-Verstraete I. 2024. Physiological role and complex regulation of O2-reducing enzymes in the obligate anaerobe Clostridioides difficile. mBio 0:e01591–24.

35. Triadafilopoulos G, Pothoulakis C, Weiss R, Giampaolo C, LaMont JT. 1989. Comparative study of *Clostridium difficile* toxin a and cholera toxin in rabbit ileum. Gastroenterology 97:1186–1192.

36. Ishida Y, Maegawa T, Kondo T, Kimura A, Iwakura Y, Nakamura S, Mukaida N. 2004. Essential Involvement of IFN-γ in Clostridium difficile Toxin A-Induced Enteritis1. The Journal of Immunology 172:3018–3025.

37. Kim H, Rhee SH, Kokkotou E, Na X, Savidge T, Moyer MP, Pothoulakis C, LaMont JT. 2005. *Clostridium difficile* Toxin A Regulates Inducible Cyclooxygenase-2 and Prostaglandin E2 Synthesis in Colonocytes via Reactive Oxygen Species and Activation of p38 MAPK*. Journal of Biological Chemistry 280:21237–21245.

38. Hryckowian AJ, Van Treuren W, Smits SA, Davis NM, Gardner JO, Bouley DM, Sonnenburg JL. 2018. Microbiota Accessible Carbohydrates Suppress Clostridium difficile Infection in a Murine Model. Nat Microbiol 3:662–669.

39. Arrieta-Ortiz ML, Immanuel SRC, Turkarslan S, Wu W-J, Girinathan BP, Worley JN, DiBenedetto N, Soutourina O, Peltier J, Dupuy B, Bry L, Baliga NS. 2021. Predictive regulatory and metabolic network models for systems analysis of Clostridioides difficile. Cell Host Microbe 29:1709–1723.e5.

40. Kelly CJ, Zheng L, Campbell EL, Saeedi B, Scholz CC, Bayless AJ, Wilson KE, Glover LE, Kominsky DJ, Magnuson A, Weir TL, Ehrentraut SF, Pickel C, Kuhn KA, Lanis JM, Nguyen V, Taylor CT, Colgan SP. 2015. Crosstalk between Microbiota-Derived Short-Chain Fatty Acids and Intestinal Epithelial HIF Augments Tissue Barrier Function. Cell Host & Microbe 17:662– 671.

41. Rivera-Chávez F, Zhang LF, Faber F, Lopez CA, Byndloss MX, Olsan EE, Xu G, Velazquez EM, Lebrilla CB, Winter SE, Bäumler AJ. 2016. Depletion of Butyrate-Producing *Clostridia* from the Gut Microbiota Drives an Aerobic Luminal Expansion of *Salmonella*. Cell Host & Microbe 19:443–454.

42. Sebaihia M, Wren BW, Mullany P, Fairweather NF, Minton N, Stabler R, Thomson NR, Roberts AP, Cerdeño-Tárraga AM, Wang H, Holden MTG, Wright A, Churcher C, Quail MA, Baker S, Bason N, Brooks K, Chillingworth T, Cronin A, Davis P, Dowd L, Fraser A, Feltwell T, Hance Z, Holroyd S, Jagels K, Moule S, Mungall K, Price C, Rabbinowitsch E, Sharp S, Simmonds M, Stevens K, Unwin L, Whithead S, Dupuy B, Dougan G, Barrell B, Parkhill J. 2006. The multidrug-resistant human pathogen Clostridium difficile has a highly mobile, mosaic genome. Nat Genet 38:779–786.

43. van Eijk E, Anvar SY, Browne HP, Leung WY, Frank J, Schmitz AM, Roberts AP, Smits WK. 2015. Complete genome sequence of the Clostridium difficile laboratory strain 630Δerm reveals differences from strain 630, including translocation of the mobile element CTn5. BMC Genomics 16:31.

44. Ng YK, Ehsaan M, Philip S, Collery MM, Janoir C, Collignon A, Cartman ST, Minton NP. 2013. Expanding the Repertoire of Gene Tools for Precise Manipulation of the Clostridium difficile Genome: Allelic Exchange Using pyrE Alleles. PLoS One 8.

45. Bouillaut L, McBride SM, Sorg JA. 2011. Genetic Manipulation of Clostridium difficile. Curr Protoc Microbiol 0 9:Unit-9A.2.

46. Olaitan AO, Dureja C, Youngblom MA, Topf MA, Shen W-J, Gonzales-Luna AJ, Deshpande A, Hevener KE, Freeman J, Wilcox MH, Palmer KL, Garey KW, Pepperell CS, Hurdle JG. 2023. Decoding a cryptic mechanism of metronidazole resistance among globally disseminated fluoroquinolone-resistant Clostridioides difficile. Nat Commun 14:4130.

47. Andrews S. FastQC A Quality Control tool for High Throughput Sequence Data. https://www.bioinformatics.babraham.ac.uk/projects/fastqc/. Retrieved 12 October 2024.

48. Bolger AM, Lohse M, Usadel B. 2014. Trimmomatic: a flexible trimmer for Illumina sequence data. Bioinformatics 30:2114.

49. Li H, Durbin R. 2009. Fast and accurate short read alignment with Burrows-Wheeler transform. Bioinformatics 25:1754–1760.

50. Li H, Handsaker B, Wysoker A, Fennell T, Ruan J, Homer N, Marth G, Abecasis G, Durbin R, 1000 Genome Project Data Processing Subgroup. 2009. The Sequence Alignment/Map format and SAMtools. Bioinformatics 25:2078–2079.

51. Bj W, T A, T S, M P, A A, S S, Ca C, Q Z, J W, Sk Y, Am E. 2014. Pilon: an integrated tool for comprehensive microbial variant detection and genome assembly improvement. PloS one 9.

52. García-Alcalde F, Okonechnikov K, Carbonell J, Cruz LM, Götz S, Tarazona S, Dopazo J, Meyer TF, Conesa A. 2012. Qualimap: evaluating next-generation sequencing alignment data. Bioinformatics 28:2678–2679.

53. Page AJ, Taylor B, Delaney AJ, Soares J, Seemann T, Keane JA, Harris SR. 2016. SNP-sites: rapid efficient extraction of SNPs from multi-FASTA alignments. Microbial Genomics 2:e000056.

54. Pensinger DA, Dobrila HA, Stevenson DM, Hryckowian ND, Amador-Noguez D, Hryckowian AJ. 2024. Exogenous butyrate inhibits butyrogenic metabolism and alters virulence phenotypes in Clostridioides difficile. mBio 15:e02535–23.

55. bcl2fastq: A proprietary Illumina software for the conversion of bcl files to basecalls.

56. Kim D, Paggi JM, Park C, Bennett C, Salzberg SL. 2019. Graph-based genome alignment and genotyping with HISAT2 and HISAT-genotype. Nat Biotechnol 37:907–915.

57. Liao, Y L, Smyth G, Shi, W. 2014. featureCounts: an efficient general purpose program for assigning sequence reads to genomic features. Bioinformatics (Oxford, England) 30.

58. R Core Team. 2020. R: A language and environment for statistical computing. R Foundation for Statistical Computing Vienna, Austria.

59. Robinson MD, McCarthy DJ, Smyth GK. 2010. edgeR: a Bioconductor package for differential expression analysis of digital gene expression data. Bioinformatics 26:139–140.

60. Schmittgen TD, Livak KJ. 2008. Analyzing real-time PCR data by the comparative CT method. 6. Nat Protoc 3:1101–1108.

